# How do body size and habitat fragmentation influence extinction in lizards? A long-term case study on artificial islands in the Brazilian Cerrado

**DOI:** 10.1101/2024.11.05.622112

**Authors:** Rogério B. Miranda, Karen C. Abbott, Ryan A. Martin, Reuber A. Brandão

## Abstract

Habitat fragmentation is known to cause extinctions and species turnover, but the factors that allow some species to persist while others become locally extinct are not well understood. Landscape flooding following the construction of hydroelectric dams causes a particularly dramatic form of fragmentation disturbance, where former terrestrial habitats become aquatic and former hilltops become land-bridge islands. As such, reservoir land-bridge islands have become a successful model for accessing the impacts of fragmentation on biodiversity. We used the lizard community, to assess species’ sensitivity to habitat structural change during land-bridge island formation. We monitored the lizard community for 23 years before, during, and after the flooding of the Serra da Mesa Dam reservoir. Over the course of our study, the diversity of the lizard community on land-bridge islands and mainland sites along the shores of the newly formed reservoir declined from 19 to six species. We found that in Serra da Mesa islands, lizards with large body sizes (e.g., Teiidae and Tropiduridae) decreased in abundance along the flooding process, thereby increasing their extinction risk. In contrast, we found a high abundance of small-bodied lizards (Gekkonidae, Gymnophthalmidae, Scincidae, and Sphaerodactylidae) on Serra da Mesa islands. Richness on the islands declined dramatically, resulting in communities currently with one highly abundant species, *Gymnodactylus amarali*. For the sake of biodiversity conservation, island or fragment sizes must be considered for maintaining a reasonable number of species and our characterization of the local extinction patterns may provide relevant information to mitigate wildlife depletion due to habitat fragmentation.

## 1 Introduction

The most concerning effect of habitat destruction is biodiversity decline. Species disappear at different points of habitat loss as each species requires a minimum amount of habitat to persist in a landscape (Fahrig 2002). Reduction of continuous habitat into smaller, scattered patches within a dissimilar matrix is known as fragmentation, one of the most common causes of habitat degradation (Crooks et al. 2017). Both biotic and abiotic factors can promote natural patchiness in the landscape and anthropogenic alterations also act to intensify habitat fragmentation worldwide (Crooks et al. 2017). A common cause of fragmentation in hyper-diverse tropical developing countries is habitat insularization promoted by river damming (Palmeirim et al. 2018). Due to an extreme level of disturbance along archipelagic landscape formation, communities face extirpation and species turnover, resulting in radical local diversity changes (Cosson et al. 1999; Gibson et al. 2013; Palmeirim et al. 2021).

Species loss during the formation of land-bridge islands after river damming is predicted by theoretical models (Simberloff 1974; Wilcox 1978; Karr 1982) and reported in empirical studies (Jones et al 2016; Palmeirim et al. 2017), but which species will go extinct remains an open question. Compared with the rarer species, the most common rodent species in a continuous tropical forest in Thailand showed the highest probability of being extirpated after the flooding of a hydroelectric dam reservoir (Lynam 1997). This result suggests that initial abundance is not a good predictor of persistence time on newly formed islands. The higher post-flooding abundance of initially rarer rodent species on the islands was considered an effect of predator decline (Lynam 1997).

Reservoir land-bridge islands have served as a successful model for assessing the impacts of island formation on isolated communities belonging to different taxa, such as mammals (Meyer & Kalko 2008; Palmeirim et al. 2018, 2020), amphibians (Brandão & Araujo 2008), birds (Yu et al. 2012), and trees (Terborgh et al. 2006). Mammals and birds are the taxa mostly widely used to assess a reservoir’s ecological impact (Jones et al, 2016; Palmeirim et al. 2017). Only recently, lizards have been studied in this context (e.g., Palmeirim et al. 2017, 2021; Miranda et al. 2023). Lizards belong to a very ecologically diverse group, are ectotherms, present low dispersal abilities, occupy a variety of trophic niches, and are highly susceptible to habitat structure alterations (Ávila-Pires 1995; Gainsbury & Colli 2019; Palmeirim et al. 2021). In addition, they play an important part in ecosystems, acting as prey, predator, and seed dispersers (Terborgh et al. 2001). All these features make lizards remarkable bioindicators (Gainsbury & Colli 2019) for extinction studies. The ecological diversity of lizards also makes them ideal for identifying species characteristics that account for permanence or extirpation in a fragmented area, which is crucial for effective conservation planning (Wang et al. 2009; Palmeirim et al. 2017).

The overarching goal of this study is to use data collected from over 20 years of monitoring in Serra da Mesa to understand the ecological drivers of local lizard extinction due to flooding. Herein, we assess ecomorphological data of lizard species that were present before the flooding of the Serra da Mesa Dam area and analyze the extirpation patterns that followed. To that end, we analyzed two different datasets: one composed of morphological data aimed at explaining the variation in lizard species persistence on these artificial islands; and another assembling the abiotic island traits of size and isolation. Specifically, we asked: 1) what are the patterns of species diversity loss during the flooding period?; 2) are extirpation rates different between islands and mainland sites?; 3) do island size and isolation affect extirpation on islands?; and 4) can body size predict which lizard species will be extirpated on island and mainland sites? Our hypothesis is that not all lizard species are impacted by habitat fragmentation in the same way. Anecdotally, after many years of monitoring, we noticed a dramatic decline in larger and more common lizards on the island sites, but this pattern has not been tested formally. We predicted that body size could explain the current configuration of communities and that area loss, habitat impoverishment, and isolation – as the degree of isolation from other islands and the mainland affects the colonization rates (MacArthur & Wilson 1967) – also shape the patterns of species diversity decline on islands.

## 2 Material and Methods

### 2.1 Study area

The Serra da Mesa dam is located in the northern State of Goiás, Central Brazil, in the core of the Cerrado biome (48° 20’W; 13° 51’S). The Serra da Mesa lake covers 178 km^2^ and was formed after the damming of the Tocantins River, a large affluent of the Amazon River. The valley flooding isolated the highest hilltops, forming nearly 280 islands. Most of these islands (about 80%) are less than 3 ha, but some are larger than 1000 ha. The flooding process began in October 1996 and water depths continued to increase until around January 1999.

Before the flooding, eight hills (predicted to be future islands) were chosen for monitoring, and after the end of the flooding process, five of these islands remained: I-34, I-35, I-37, I-38, and IX (Fig. 1). The islands vary in size from 3 to 35 ha. The choice of these areas was based on vegetation type, level of human disturbance (fire, cattle grazing, logging), and accessibility. Due to fluctuations in water level, island IX reconnected with the mainland. Five additional sites on the mainland (M1-M5, Fig. 1) were monitored as well for comparison with the island sites.

**Figure 1.**
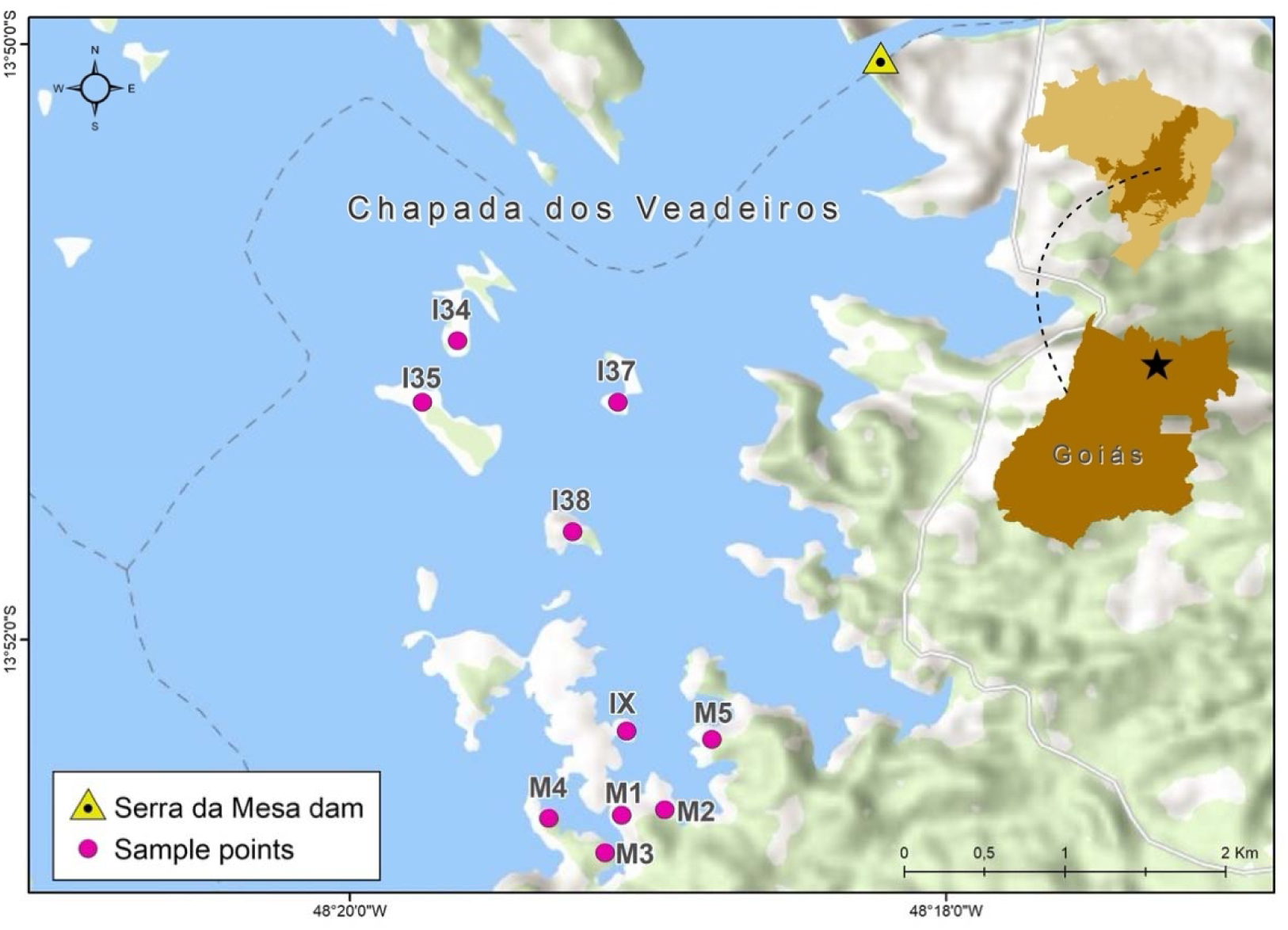
Map of the Serra da Mesa dam and lake in Goiás State, Brazil, showing the monitoring sites. Insets show a map of Brazil (upper) with the Cerrado region shaded, and the state of Goiás (lower) that is centrally located in this region.

### 2.2 Species survey and monitoring

Lizard richness, abundance, and species presence were obtained through visual survey, pitfall trap captures, and sampling after controlled burns. The visual survey was carried out from July to October 1996, followed by bimonthly pitfall trapping until January 1999. We installed 20 L buckets in grids of 15 traps, and each trap grid covered an area of 72 m^2^. We left the traps open for 13 days, monthly. A total of 315 traps (21 grids) were installed in the studied hills. We took measurements, marked by toe clipping, and released all the captured lizards. Lizards observed outside the traps were identified with habitat and activity time recorded.

In 2001, 2011, and 2019, we sampled lizards in ten 50mx50m square plots enclosed by plastic fences. The change in methodology was necessary because the rocky soil at the hilltops and the reduced sample area after flooding made digging burrows for the 20 L pitfall buckets nearly impossible. This methodology, which we named fire squares, was used in previous studies (e.g., Brandão 2002; Amorim 2017) with satisfactory efficiency. We delimited the area with tapelines and enclosed the square using plastic canvas stretched between wooden stakes. After that, we made a 3-meter-wide firebreak around the fenced areas and burned the vegetation to expose all possible shelters used by lizards. After removing the vegetation, we carefully searched for lizards, checking all available shelters such as holes, trunk logs, rock crevices, and termite mounds. We controlled for the capture of all lizards in the enclosure using rarefaction curves.

After two hours of sampling without any lizards recorded, we considered that all lizards in the square were captured. There were no dead or wounded lizards due to the use of this sampling method. All lizards captured were measured, weighed, euthanized with lidocaine injection in the coelom, fixed in formalin, and posteriorly transferred to 70% alcohol for preservation. All animals were housed at Coleção Herpetológica da Universidade de Brasília (CHUNB).

### 2.3 Morphological data

Snout-vent length (SVL) and mass were measured for most animals collected during the surveys described above, though data were missing for some of the species. Using information from Meiri (2024) and personal communication, we filled in the missing data. A detailed explanation is presented in the supplementary material. We acknowledge the fact that mixing field measurements and literature data is not ideal; however, it was necessary because neither data source was complete. Some species, particularly the most common ones, lack natural history data in the scientific literature, and even studies with broad datasets (e.g. Meiri 2024) dealt with many gaps in their metadata. Thus, only by combining our own field measurements with published data, we were able to assemble a reasonably complete morphological dataset. Note, however, that we did not include *Hoplocercus spinosus* and *Tropidurus torquatus* in our analyses because SVL and mass data were not available, although they were reported at our study sites.

### 2.4 Statistical analyses

#### 2.4.1 What are the patterns of species diversity loss during the flooding period?

To answer this question, we calculated richness, abundance, and beta diversity at each census point between 1996 and 1999 to evaluate changes in the species composition and the monitoring sites during the flooding process. For this period, we estimated lizard richness and abundance using mark-recapture in pitfall traps and visual surveys. Beta diversity was measured using the Whittaker index (Whittaker 1972) to measure differences in local species composition between pairs of sites using the equation BW=(*S*/*a*)-1, where *S* is the total number of species found at either site, and *a* is the mean richness within the sites. The beta diversity index values range from 0 to 1, where values nearest to 1 show a larger turnover.

Differences in beta diversity, richness, and abundance on islands before, during, and after the flooding were tested with the Mann-Whitney test. We performed a simple linear regression analysis to test the relationship between species abundance and time (data bimonthly grouped, 1996-1999) to assess abundance changes in the local lizard community along the flooding process.

#### 2.4.2 Are extirpation rates different between islands and mainland sites?

We constructed a generalized linear mixed-effects model to evaluate if lizards on island sites had greater risks of extirpation than lizards on mainland sites across the duration of our survey period (1996-2019). Our response variable was the proportion of lizard species at each site at each census point, out of the 17 species pool, excluding *Hoplocercus spinosus* and *Tropidurus torquatus*. Our fixed effects were site type (island or mainland), census year as the number of years post flooding, and the interaction between year and site type. We used a beta error distribution, included census site as a random intercept, and included an auto-regressive correlation structure to account for temporal nonindependence among successive census points (Box et al. 1994; Pinheiro and Bates 2000). We fit this model using the glmmTMB function from the {glmmTMB} library in R (Brooks et al. 2017), and tested significance of the fixed effects using analysis of deviance, implemented by the ANOVA function in the {car} library (Weisberg, 2019).

#### 2.4.3 Does island size and isolation affect extirpation on islands?

For this question, we constructed a generalized linear mixed-effects mode to determine if island size or isolation (distance from nearest other island or the mainland shore) influenced the extirpation of lizard species over time. The response variable was the proportion of lizard species on each island at each census point out of the total starting species pool. We included island size and distance as fixed effects, both natural log-transformed along with census year, and the two-way interaction between year and the island variables. We did not include a 3-way interaction as this model would not converge. We used a beta error distribution and included census site as a random intercept. We fit this model using the glmmTMB function in the eponymous library and tested the significance of the fixed effects using type 3 analysis of deviance, implemented by the ANOVA function in the {car} library.

#### 2.4.4 Can body size predict which lizard species will be extirpated on island and mainland sites?

Lastly, we constructed mixed-effect Cox proportional hazards models using all species for whom we had SVL and mass data to ask if larger lizard species had greater risks of extirpation over time since the reservoir’s construction. Because a model including site type (island or mainland), lizard size, and their interaction did not meet the assumptions of proportional hazards, we ran separate models for extirpation risk on the island sites and the mainland sites. For both island and mainland sites, we ran separate models for body mass (g) and SVL (mm), as they were highly correlated (r = 0.96). Our response variable was the retention or loss (coded as 0 and 1) of each lizard species at each site, at each census point. We log-transformed both measures of body size due to the large range of size values in the dataset. We included a random intercept of site and a separate random intercept to account for phylogenetic relatedness by nesting species within genus, and species and genus within family. We fit these models with the *coxme* function from the eponymous library (Therneau, 2022) and tested the significance of the fixed effects using type 2 analysis of deviance, implemented by the ANOVA function in the {car} library. All statistical analyses were done in R Console software version 4.2.3 (R Development Core Team, 2023), except for the differences in beta diversity, richness, and abundance, which were performed using the softwares SYSTAT 7.0 and SAS.

## 3 Results

### 3.1 What are the patterns of species diversity loss during the flooding period?

We recorded 19 lizard species in the sampled area before and during the flooding, distributed among 10 families: Teiidae (4 species), Gymnophtalmidae (3), Tropiduridae (3), Anolidae (2), Scincidae (2), Hoplocercidae (1), Iguanidae (1), Phyllodactylidae (1), Polychrotidae (1), and Sphaerodactylidae (1) (Table 1). We found 15 species at the hilltops and 17 within the area that would be flooded. Two species (*Tropidurus montanus* and *Anolis meridionalis*) were recorded only at the hilltops. Four species were only observed in the valley-flooded habitats prior to the flooding, mainly in gallery forests and adjacent open Cerrado (*Tupinambis quadrilineatus, Tropidurus torquatus, Hoplocercus spinosus*, and *Polychrus acutirostris*), though some of these were later seen on hilltop islands.

**Table 1.**
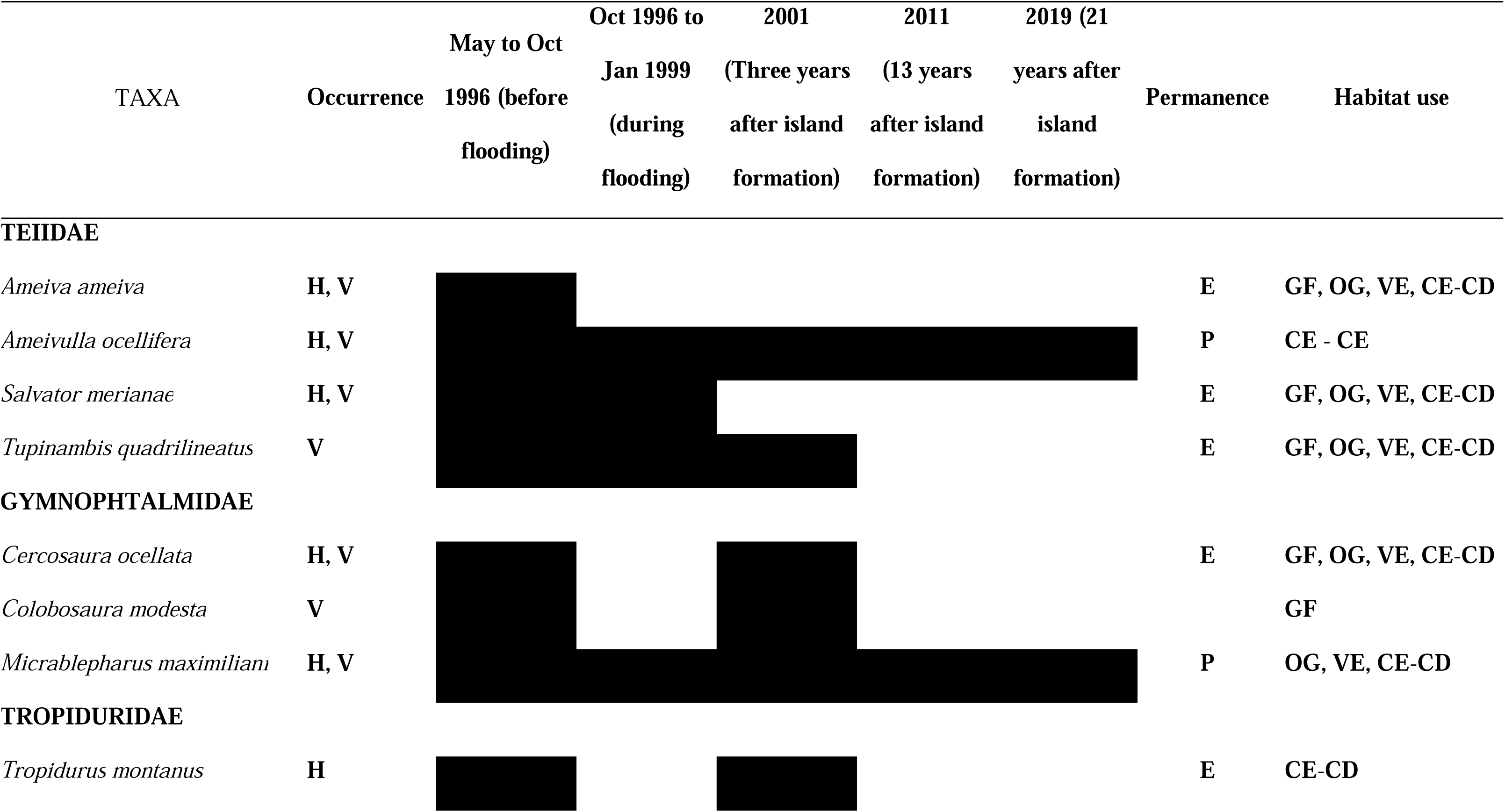
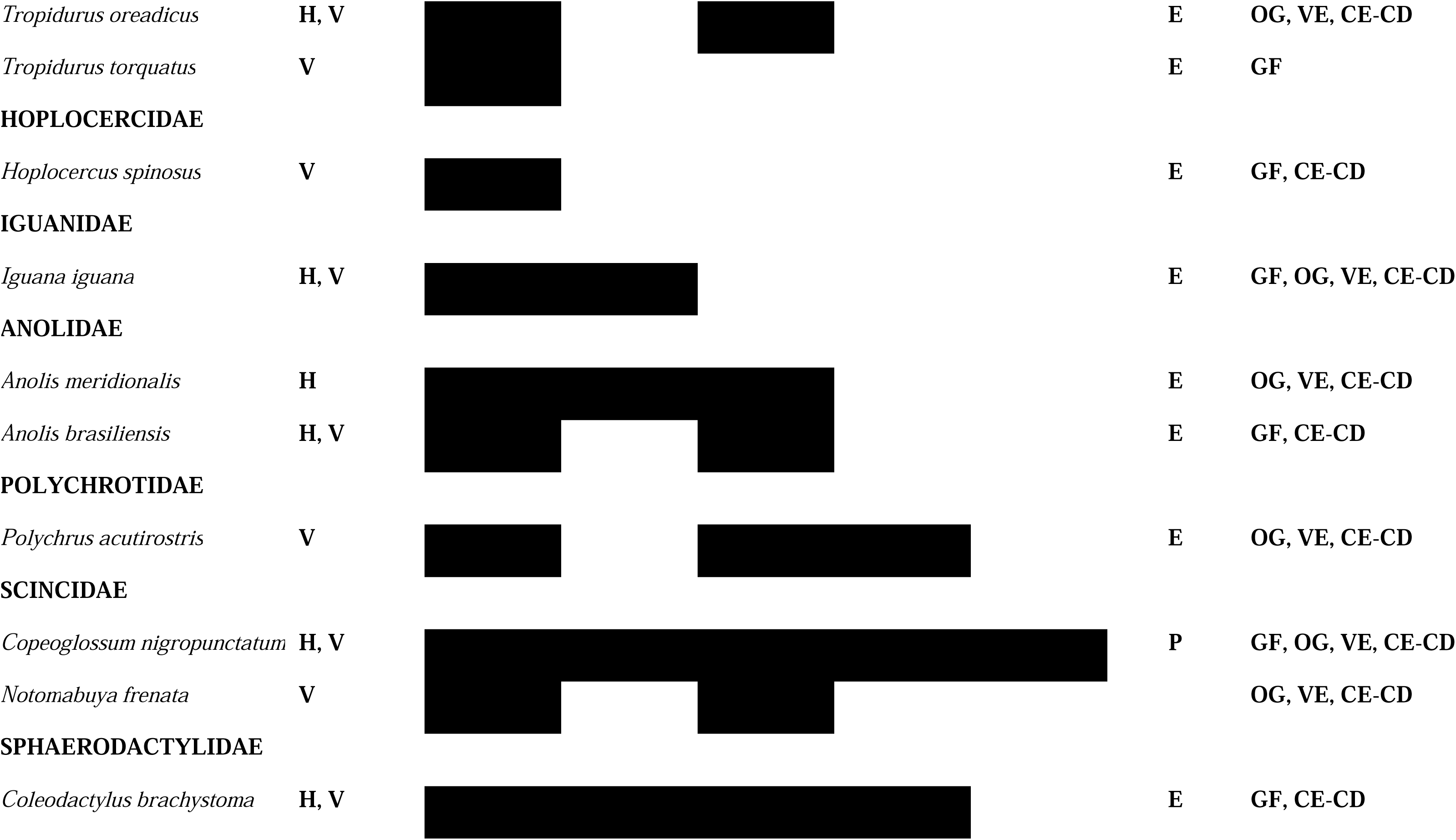
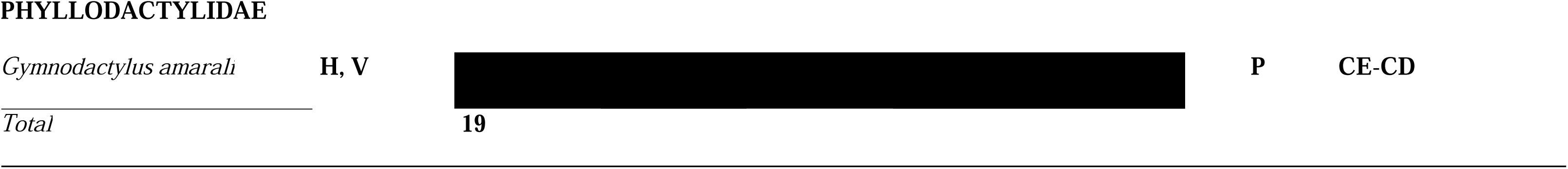
Lizard species in the study area distributed by sampling period on island habitats. H – Species recorded at hilltops; V – Species recorded at valleys (that became flooded); P – Species recorded on islands in 2019; E – Species extirpated. The habitats used by the lizards are gallery forest (GF), open grassland (OG) veredas (VE), and cerrado-cerradão (CE-CD).

Within the sampled area, we detected high lizard richness and seven species endemic to the Cerrado biome (Colli et al. 2002). The data show that the early saurofauna from Serra da Mesa was predominantly composed of heliophile and habitat-generalist species.

The beta diversity was calculated for all combinations of island pairs (mainland excluded) before flooding, island pairs after the flooding, and before versus after flooding for the same island. The mean beta diversity between hilltops before the flooding (1996) was 0.434 ± 0.148, whereas the mean beta diversity between the same hilltops that became islands after the complete flooding (1999) was 0.524 ± 0.181, which shows a tendency of greater dissimilarity after the flooding. This difference, however, was not statistically significant (U_(1,56)_= 276.00; p= 0.057).

Between 1996 and 1999 we captured 485 lizards from 13 different species using pitfall traps. The regression between the abundance of each captured species against time (bimonthly), showed that the Teiidae lizards, *Ameiva ameiva* (slope = −0.530, p = 0.035) and *Ameivula ocellifera* (slope = −0.498, p= 0.050), and the Tropiduridae lizards, *Tropidurus oreadicus* (slope = −0.668, p = 0.005) and *Tropidurus montanus* (slope = −0.526, p = 0.016), decreased their abundance between July 1996 and January 1999 (Fig. 2). The large *A. ameiva* was last recorded in 1998 and it was considered locally extinct on the islands.

**Figure 2.**
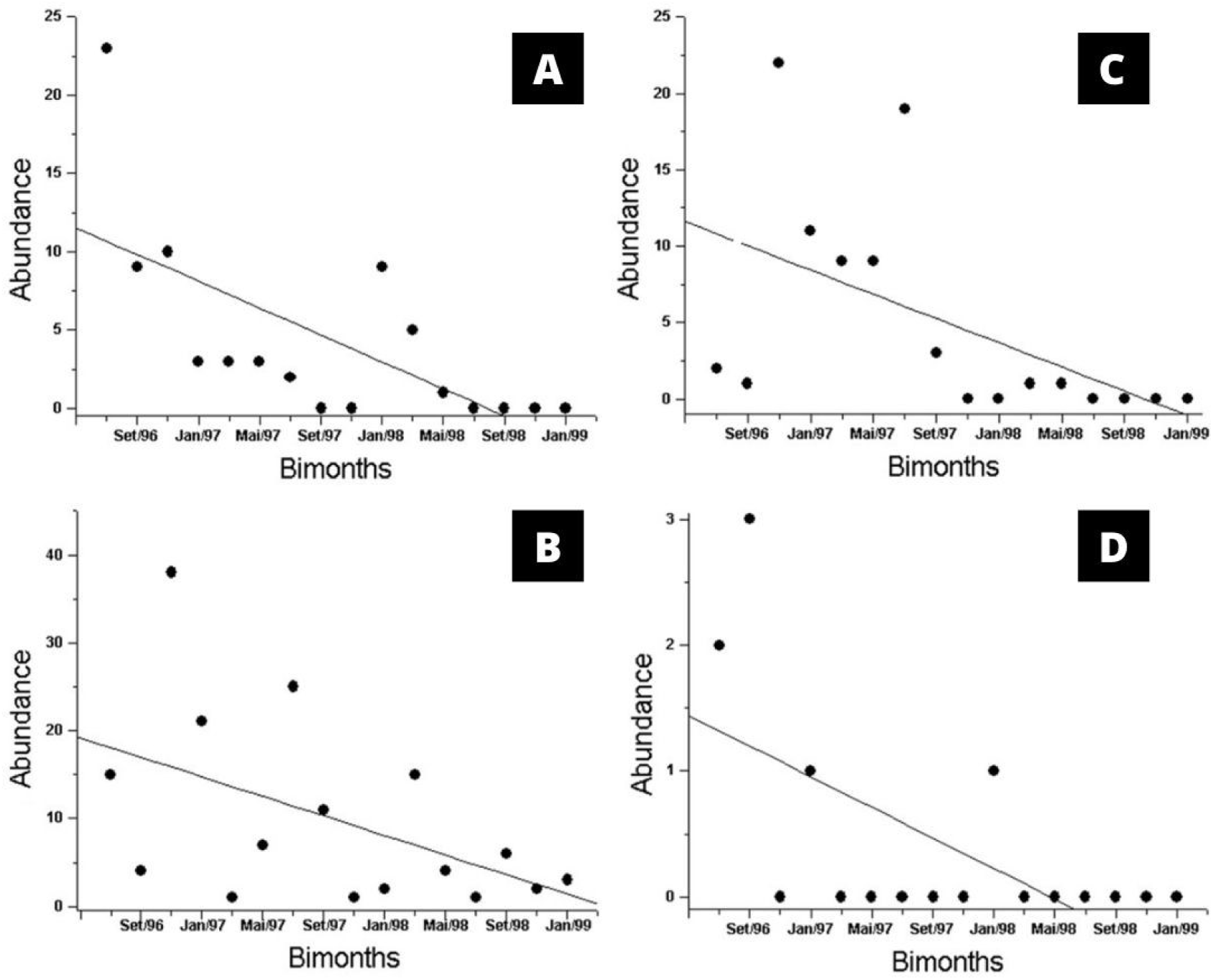
Regressions between the abundance of the four largest lizards in our region and time (bimonthly). A. *Ameiva ameiva*; B. *Ameivulla ocellifera*; C. *Tropidurus oreadicus*; D. *Tropidurus montanus*.

*Ameiva ameiva, Ameivula ocellifera,* and *Tropidurus oreadicus* were the most abundant lizards at the hilltops before the flooding. These species and *T. montanus* were the largest lizards from the studied community (excluding tegus and iguanas, not captured by the pitfall traps). Abundance declines of *A. ameiva* and *T. oreadicus* were also detected at the mainland sites, during the reservoir formation (Pavan 2001). The remaining nine lizard species are smaller, discretive, and rarer, with more restricted geographic distribution in some cases. Interestingly, these species did not show a significant drop in abundance during island formation (Fig. 3).

**Figure 3.**
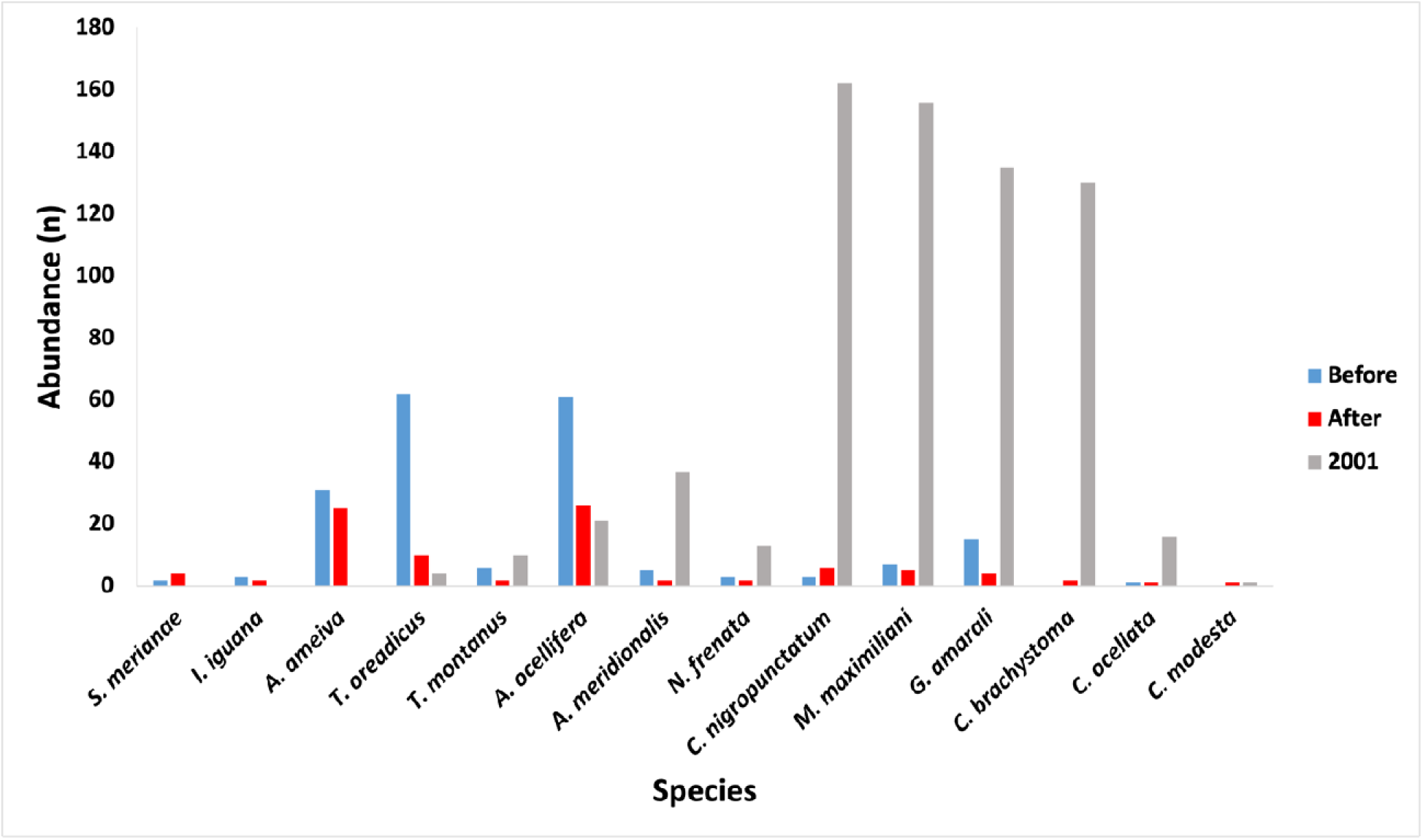
Abundance of lizard species at Serra da Mesa, comparing the first 12 months (1996-1997, before the flooding), the last 12 months (1998-1999, after the flooding), and the 2001 sampling. Abundance in number of individuals (n). Only data for the 14 species reported during the flooding or on islands after reservoir formation are shown. The species are arranged along the x-axis in approximate size order, from larger to smaller.

**Figure 4.**
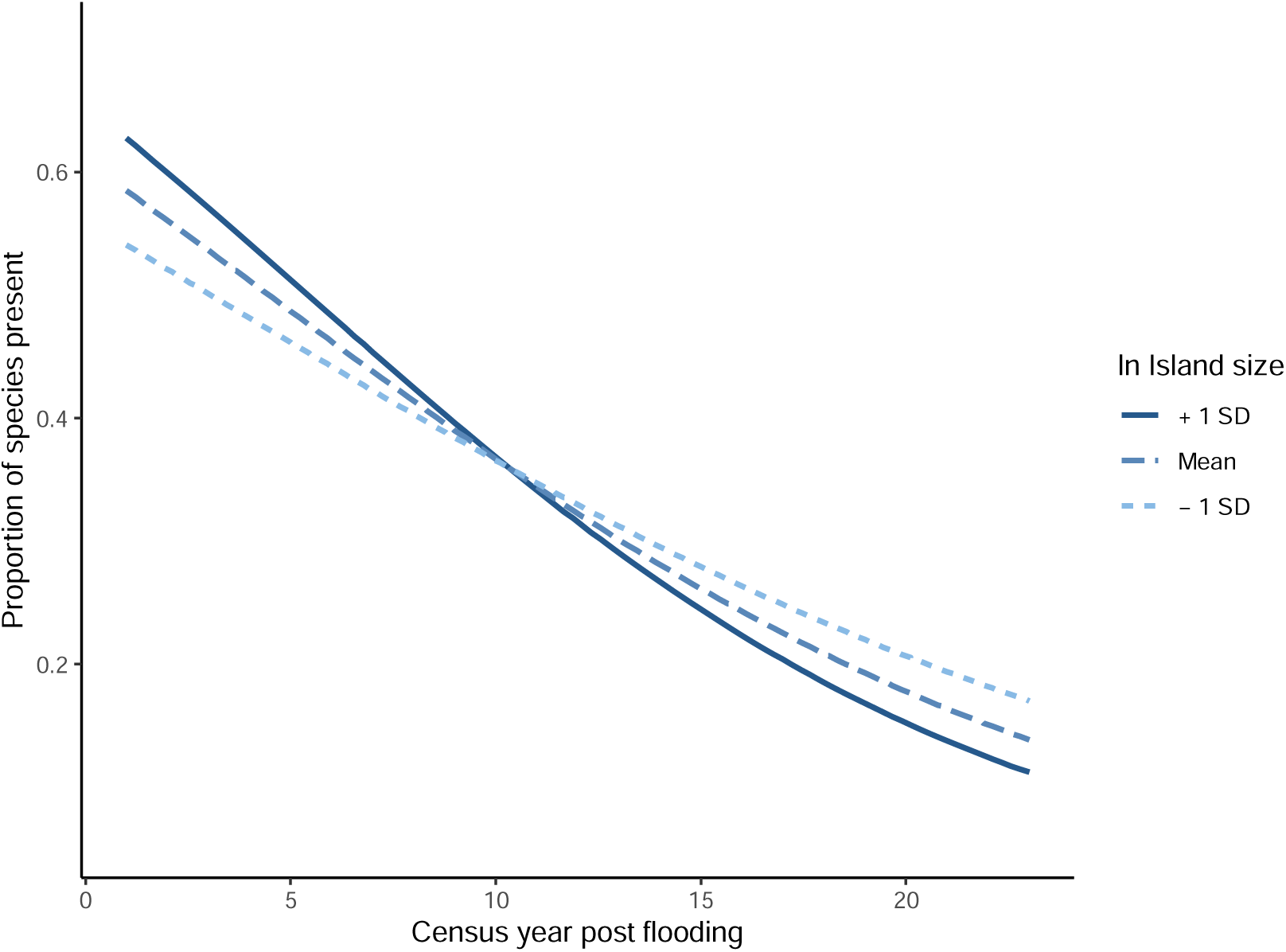
Visualization of the significant year by island size interaction, predicted from a generalized mixed model (mean relationship shown by the dark-blue dashed curve ± 1 SD in the natural log of island size). Larger islands tended to have greater species richness initially following island flooding but lower species richness by the final census. Island size, natural log transformed, from smaller to larger is indicated by point color, from lighter to darker shades.

Nearly all species reported showed abundance decline at the end of the reservoir filling, even those that were smaller, more discretive, and rarer, with more restricted geographic distributions. Interestingly, these smaller species presented an impressive recovery by the 2001 sampling (Fig. 3). During the sampling in 2001, we found 686 lizards belonging to 12 species on the islands, whereas 109 specimens were recorded on the mainland. The number of lizards found on islands was six times that on the mainland. Four lizard species were dominant in the island communities: *Copeoglossum nigropunctatum* (162 individuals or 23.6% of all lizards found), *Micrablepharus maximiliani* (156; 22.75%), *Gymnodactylus amarali* (135; 19.68%), and *Coleodactylus brachystoma* (130; 18.95%). These four species (one-third of species found) accounted for 73.33% of all sampled individuals. These lizards were not as abundant in the samples taken during the flooding. In contrast, the larger, dominant species during the flooding between 1996 and 1999 (*Ameiva ameiva, Ameivulla ocellifera,* and *Tropidurus oreadicus*) were rare three years after the complete flooding (the 2001 sample), representing, respectively, 0% (locally extinct), 3%, and 0.6% of the total abundance in the isolated communities, a severe abundance decline. Similarly to *A. ameiva*, *Salvator merianae* and *Iguana iguana*, two other large-bodied species, were never recorded on the islands after the reservoir filling was over.

In 2011, we recorded 284 individuals from 12 species, spread among eight families (Anolidae, Gymnophtalmidae, Phyllodactylidae, Polychrotidae, Sphareodactylidae, Scincidae, Teiidae, and Tropiduridae). On the mainland, 160 specimens were collected, while on the islands we captured 124 individuals from eight species. *G. amarali* was the most abundant lizard on the islands, corresponding to 44.6% of all individuals found. The only lizard species found exclusively on the islands in 2011 was *Polychrus acutirostris,* wherea*s Anolis brasiliensis, A. meridionalis, Tropidurus montanus,* and *Salvator merianae* were restricted to the mainland.

In 2019, we recorded 24 individuals distributed across only six species. We found 15 individuals belonging to five species on the islands, with *G. amarali* corresponding to 40% of the total. *Colobosaura modesta* was only recorded on the mainland.

### 3.2 Does extirpation risk differ between mainland and island sites?

The proportion of lizard species out of the total pre-flood species pool fell across the duration of the study (slope = −0.098 ± 0.009, ***χ***^2^ = 80.1, *p* < 0.0001). The mean proportion of species present on mainland and island sites was not significantly different (island: 0.376 ± 0.093; mainland: 0.319 ± 0.027, ***χ***^2^ = 2.22, *p* = 0.136) averaged across the monitoring time. While the rate of species extirpation tended to be greater on islands compared to mainland sites (island slope: −0.099 ± 0.011; mainland slope: −0.073 ± 0.011), this interaction between site type and census year was non-significant ***χ***^2^ = 2.91, *p =* 0.088).

### 3.3 Does island size or isolation influence the risk of lizard extirpation?

There was no evidence that island isolation (***χ***^2^ = 0.14, *p* = 0.707) or the interaction between island isolation and year influenced lizard extirpation (***χ***^2^ = 0.832, *p* = 0.362). There was, however, a significant interaction between island size and census year (slope = −0.016 ± 0.007, ***χ***^2^ = 5.01, *p* = 0.025). Plotting this interaction suggests that larger islands had greater species diversity at the beginning of the study, consistent with the Theory of Island Biogeography (MacArthur & Wilson 1963; MacArthur & Wilson 1967) but had lower lizard diversity at later census points (Fig. 3).

### 3.4 Can body size predict which lizard species will be extirpated on island and mainland sites?

We found that larger lizard species had higher risks of extirpation on island sites. This was true for both measures of size (Fig. 5A, B). Specifically, there was a 77% increase in extirpation risk for a 10% increase in SVL (hazard ratio = 1.77 ± 0.269, ***χ***^2^ = 4.56, p = 0.0327) and a 20% increase in extirpation risk for a 10% increase in mass (hazard ratio = 1.20, ***χ***^2^ = 4.898 ± 0.0839, p = 0.0269). In contrast, neither SVL nor mass predicted extirpation risk for the mainland sites (ln svl: hazard ratio = 1.11 ± 0.0689, ***χ***^2^ = 2.15, p = 0.143; ln mass: hazard ratio = 1.03 ± 0.0219, ***χ***^2^ = 2.19, p = 0.139), although the trends remained the same with somewhat higher but statistically non-significant risks associated with species of larger size (Fig 5C, D).

**Figure 5.**
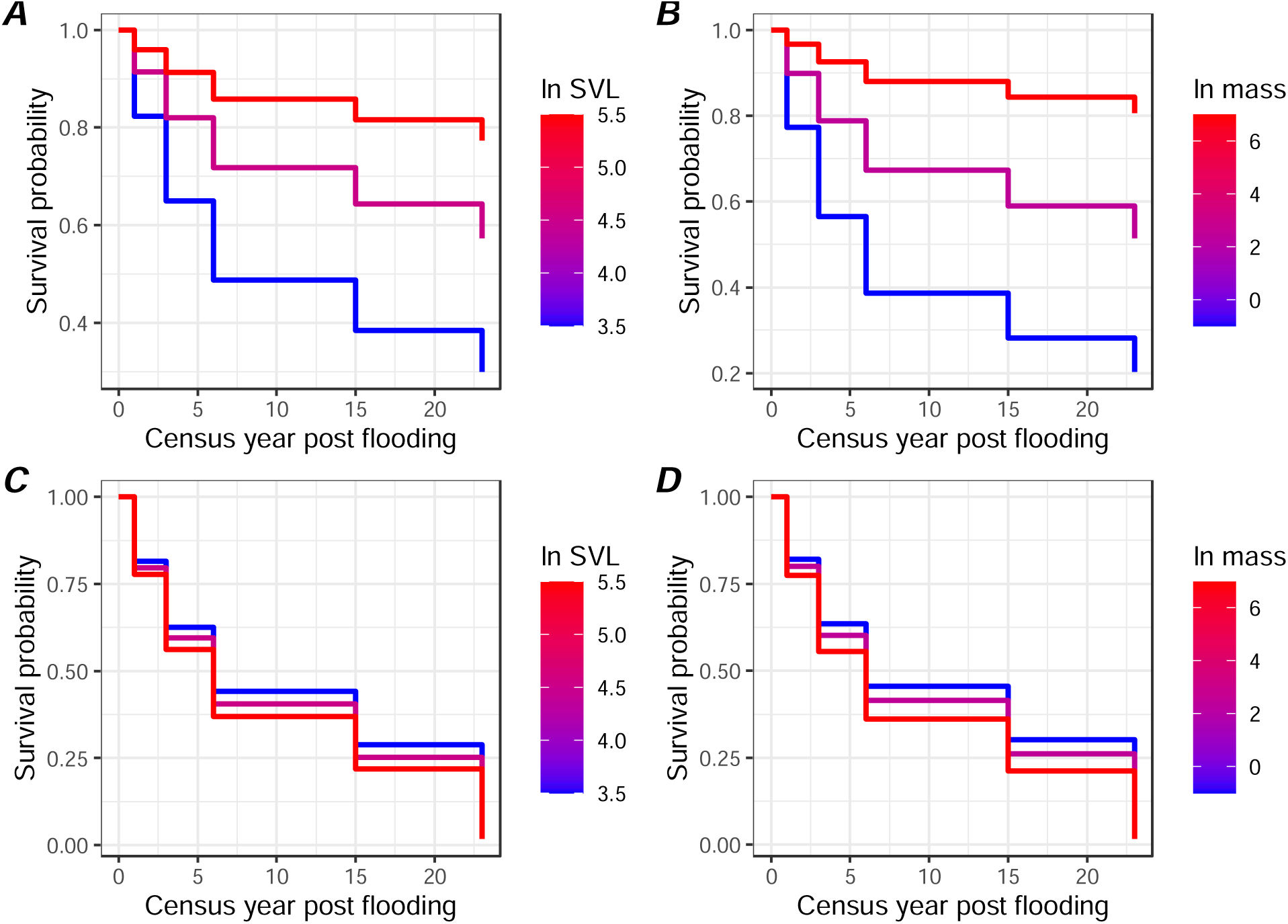
Visualization of survival probabilities across the post flooding census years for island (A, B) and mainland (C, D) lizard species in relation to mean species body size. As there are currently no existing methods to visualize survival curves from mixed-effect Cox proportional hazards models, the depicted survival curves are from models without the random effects of site or taxonomy fit using the coxph function in the {survival} library (Therneau 2024). Consequently, these survival curves should be interpreted only as representative of the mixed effects Cox models.

## 4 Discussion

The results confirm our hypothesis that habitat fragmentation affects different lizard species in different ways, with larger-bodied species being more at risk from insularization. The decline in abundance and species richness from 1996 until 2019 is severe and evident. The local lizard community dropped from 19 species and 10 families to only six species and five families. Larger, diurnal, and habitat-generalist lizards became absent on islands while smaller, discretive, and more habitat-specialist species became dominant, which suggests a pattern in the local extinction event. Although a decrease in the beta diversity between island pairs was expected due to the loss of more sensitive species, the mean beta diversity on islands showed an increasing trend after flooding, suggesting more idiosyncratic changes in the species composition during the flooding process. One reason for this increase could be the natural migratory movement of species from the valley to the higher grounds, prompted by the rise of the waters during the dam filling. This is plausible, as species initially found only in the valleys were indeed later captured on islands. Still, the contribution of any such movement to beta diversity trends was not directly measured.

Gainsbury & Colli (2003) report that extinctions in Amazonian Cerrado enclaves were stochastic consequences of historical factors. In contrast, several other studies report greater persistence of more generalist and abundant species following habitat fragmentation in forests (Smith et al. 1996; Sarre et al. 1995; Sarre 1998), and on islands and edges of hydroelectric dam lakes (Terborgh et al. 1997; Cosson et al. 1999; Palmeirim et al. 2017). However, our results detected the permanence of smaller, specialist, and initially less common lizards, which differs explicitly from what has been previously observed in similar systems, as described above. The species that declined during the Serra da Mesa flooding (from 1996 to 1999) are easily found in Cerrado remnants located around the city of Brasília (Colli 1992; Brandão & Araujo 2001; Colli et al. 2002), suggesting that the island formation in Serra da Mesa experienced different community (dis)assembly processes when compared to other fragmentation events reported for the Cerrado. Species loss thus appears to be context-dependent, shaped perhaps by the type of fragmentation (urbanization versus flooding) or other aspects of a site’s history.

Fragment size, connectivity, and distance from the nearest fragment influenced the lizard richness in the fragments investigated by Smith et al. (1996), which is expected according to the Island Biogeography model (MacArthur & Wilson 1967). We, on the other hand, saw no effect of distance in our study. Although we did not find a significant difference in extirpation rates between islands and mainland sites, our analysis suggests that larger islands had a greater number of species at the beginning of the monitoring but declined sharply in diversity later. At first, all islands experienced a loss in diversity; however, a recovery of the smaller species on the islands was reported in the 2001 census, as shown in Figure 3. It is possible that a general reduction in food and spatial resources during the islands’ formation could have favored smaller species, with lower energy demands (e.g., less daily food intake) and perhaps then an advantage in poorer habitats. Similarly, gekkonid and scincid lizards are the most common on ocean islands (Carlquist 1965). Thus, our study reveals that in the Serra da Mesa community, the smaller, secretive, and rarer species have the advantage of surviving when compared with the larger, generalist, and more conspicuous lizards. These results suggest a directional selection event in the lizard community due to the Serra da Mesa reservoir flooding.

During the flooding, there was a high occurrence of birds of prey, such as eagles, hawks, and falcons (Accipitridae, Falconidae) on the islands and several events of predation were observed throughout the fieldwork. Some bird species considered important reptile predators, like the red-legged seriema (*Cariama cristata*) and the guira cuckoo (*Guira guira*), became very common on the islands along the flooding process (Redford & Peters 1986; Souza et al. 2022). Raptors are visually oriented predators and, excluding owls, they are diurnal hunters (Jaksić 1983). Thus, larger, diurnal, and initially more abundant lizards, such as *A. ameiva, A. ocellifera,* and *Tropidurus spp*., are expected to be more vulnerable to these predators than species with crepuscular and nocturnal behavior – case of the Gekkonids (Vitt 1991) - with smaller size and/or discretive habits, like *C. brachystoma, G. amarali, M. maximiliani,* and *mabuids*. These “hidden lizards” also show a strong association with certain kinds of retreat sites – e.g. termite mounds, leaf-cutter ant mounds, and debris (Colli et al. 2002). The decline of species with these former features suggests another mechanism, in addition to higher energetic demands, for the biased loss of larger species.

In conclusion, our predictions that body size and habitat loss could explain the diversity decline of the Serra da Mesa’s lizard community were confirmed. Our results show that fragmentation, morphology (larger body size), and possible ecological (predation) and physiological (less daily food intake) factors determined the community structure in the Serra da Mesa land-bridge islands. Although considered a clean energy source, hydroelectric dams are also responsible for environmental destruction, causing fragmentation, degradation, and habitat impoverishment (Lees et al. 2016; Jones et al. 2016; Palmeirim et al. 2018). As the most concerning effect of habitat destruction is biodiversity decline, hydroelectric constructors, for the sake of biodiversity conservation, must consider the fact that island or fragment sizes are crucial for maintaining a reasonable number of fauna species. We highlight the role of long-term studies for a better understanding of fragmentation processes and emphasize that careful planning before the flooding initiation would be essential in order to mitigate wildlife impoverishment.

---

**Supplementary material.**
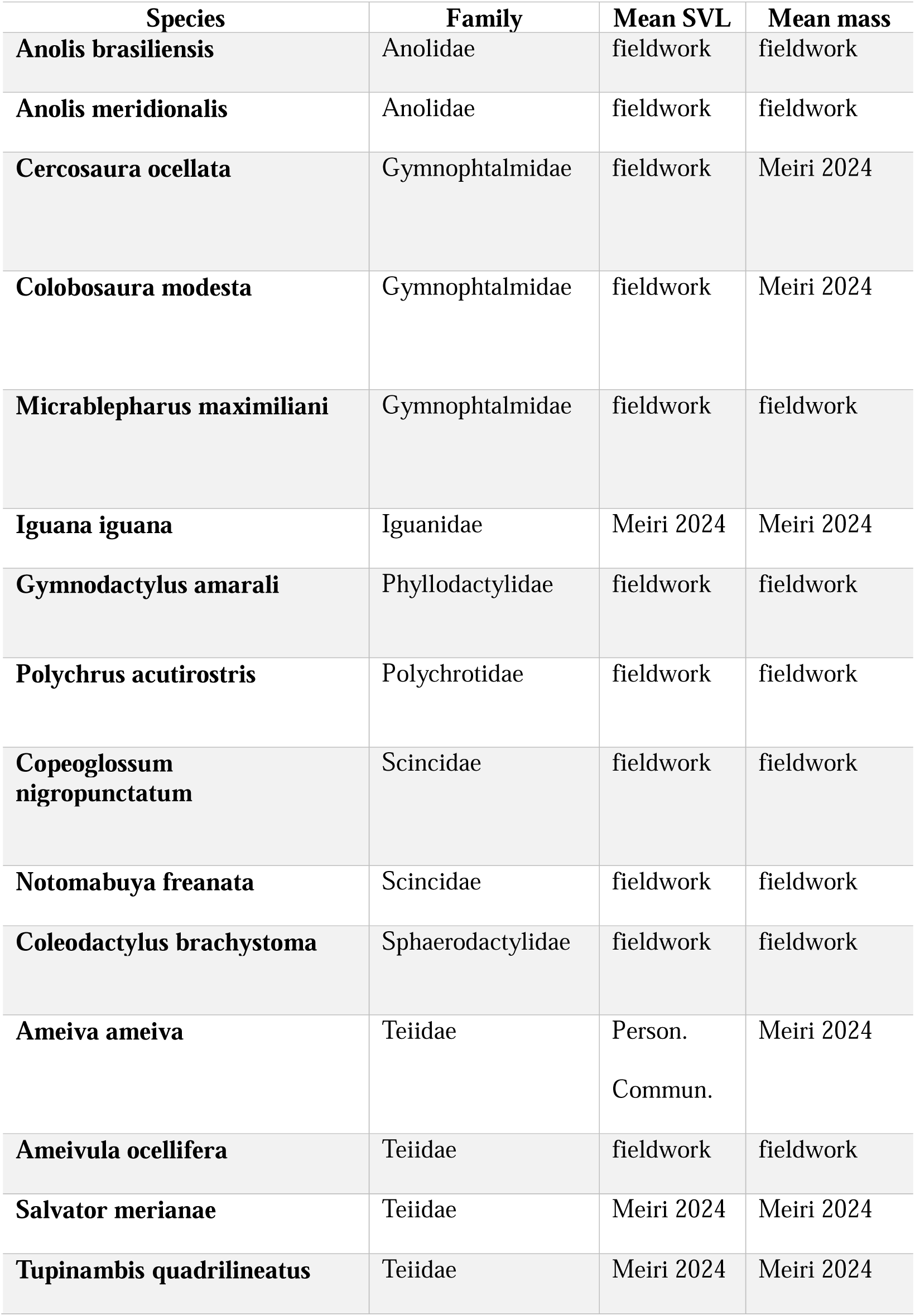
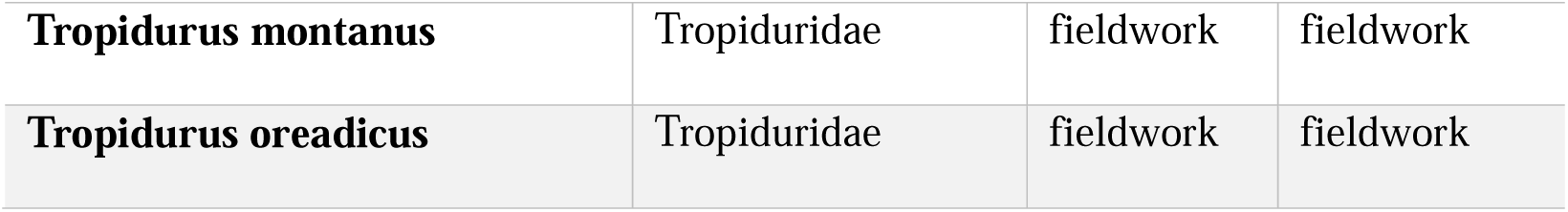
Description of data sources.

## 5 References

Amorim M., Schoener T. W., Santoro G. R. C. C., Lins A. C. R., Piovia-Scott J., Brandão R. A. (2017). Lizards on newly created islands independently and rapidly adapt in morphology and diet. Proceedings of the National Academy of Sciences, 114:8812–8816.

Adler, G. H. & Levins, R. (1994). The island syndrome in rodent populations. The Quarterly Review of Biology, 69:473–490.

Araujo. A. F. B. (1991). Structure of a white sand-dune lizard community of coastal Brazil. Revista Brasileira de Biologia, 51: 857–865.

Araujo, A. F. B. (1992). Estrutura morfométrica de comunidades de lagartos de áreas abertas do litoral Sudeste e Brasil Central. Unpublished Phd Thesis, Universidade de Campinas. 191 p.

Araujo, A. F. B. & Machado, R. B. (2000). Fragmentação de hábitats e a conservação da avifauna e herpetofauna no Cerrado do Distrito Federal. Relatório Técnico - FAPDF.

Ávila-Pires, T. C. S. D. (1995). Lizards of brazilian amazonia (Reptilia: Squamata). Zoologische verhandelingen.

Baselga, A. & Orme, C. D. L. (2012). betapart: an R package for the study of beta diversity. Methods in ecology and evolution, 3(5): 808–812.

Bierragaard Jr., R. O. & Lovejoy, T. E. (1989). Effects of forest fragmentation on Amazonian understory birds. Acta Amazonica, 19:215–241.

Box, G.E.P., Jenkins, G.M., and Reinsel G.C. (1994). Time Series Analysis: Forecasting and Control. 3rd Edition, Holden-Day.

Brandão, R. A. & Araujo, A. F. B. (2001). A herpetofauna associada às Matas de Galeria no Distrito Federal. In: Ribeiro, J. F.; Fonseca, C. E. L. & Sousa-Silva, J. C. Caracterização e Recuperação de Matas de Galeria. EMBRAPA, Planaltina. pp. 561–604.

Brandão, R. A (2002). Monitoramento das populações de lagartos no Aproveitamento Hidroelétrico de Serra da Mesa, Minaçu, GO. Thesis (Universidade de Brasília, Brasilia, Brazil).

Brandão, R.A. & Araujo, A.F. (2008). Changes in anuran species richness and abundance resulting from hydroelectric dam flooding in Central Brazil. Biotropica, 40: 263–266.

Brooks, M. E., Kristensen, K., Van Benthem, K. J., Magnusson, A., Berg, C. W., Nielsen, A.,… & Bolker, B. M. (2017). glmmTMB balances speed and flexibility among packages for zero-inflated generalized linear mixed modeling. The R journal, 9(2): 378–400.

Case, T. J. (1975). Species numbers, density compensation, and colonizing ability of lizards on islands in the Gulf of California. Ecology, 56: 3–18.

Carlquist, S. (1965). Island Life: A Natural History of the Islands of the World. Natural History Press. Garden City, New York. 451 pp.

Colli, G. R. (1991) Reproductive ecology of Ameiva ameiva (Sauria: Teiidae) in the cerrado of Central Brazil. Copeia, 1991:1002–1012.

Colli, G. R., Bastos, R. P. & Araujo, A. F. (2002). 12. The Character and Dynamics of the Cerrado Herpetofauna. In The Cerrados of Brazil (pp. 223–241). Columbia University Press.

Cosson, J. F., Ringuet, S., Classens, O., Massary, J. C. de, Dalecky, A., Villiers, J. F., Granjon, L. & Pons, J. M. (1999). Ecological changes in recent land-bridge islands in French Guiana, with emphasis on vertebrate communities. Biological Conservation, 91:213–222.

Diamond, J. M. (1972) Biogeographic kinetics: Estimation of relaxation times for avifaunas of southwest pacific islands. Proceedings of National Academy of Sciences, 69:3199–3203.

Faeth, S. H. & Connor, E. F. (1979) Supersaturated and relaxing island faunas: A critique of the species-age relationship. Journal of Biogeography, 311–316.

Gainsbury, A. M. & Colli, G. R. (2003). Lizard Assemblages from Natural Cerrado Enclaves in Southwestern Amazonia: The Role of Stochastic Extinctions and Isolation. Biotropica, 35(4): 503–519.

Gainsbury, A. M., & Colli, G. R. (2019). Phylogenetic community structure as an ecological indicator of anthropogenic disturbance for endemic lizards in a biodiversity hotspot. Ecological Indicators, 103: 766–773.

Gibson, L., Lynam, A. J., Bradshaw, C. J., He, F., Bickford, D. P., Woodruff, D. S.,… & Laurance, W. F. (2013). Near-complete extinction of native small mammal fauna 25 years after forest fragmentation. Science, 341(6153): 1508–1510.

Herrmann, N. C., Stroud, J. T., & Losos, J. B. (2020). The Evolution of ‘Ecological Release’ into the 21st Century. Trends in Ecology & Evolution.

Jaksić, F. M. (1983). The trophic structure of sympatric assemblages of diurnal and nocturnal birds of prey. American Midland Naturalist, 152–162.

Jones, I. L., Bunnefeld, N., Jump, A. S., Peres, C. A. & Dent, D. H. (2016) Extinction debt on reservoir land-bridge islands. Biological Conservation, 199: 75–83.

Karr, J. R. (1982). Population variability and extinction in the avifauna of a tropical land bridge island. Ecology, 1975–1978.

Lees, A. C., Peres, C. A., Fearnside, P. M., Schneider, M. & Zuanon, J. A. (2016) Hydropower and the future of Amazonian biodiversity. Biodiversity and conservation, 25(3): 451–466.

Lynam, A. J. (1997). Rapid decline of small mammal diversity in Monsoon evergreen forest fragments in Thailand. In: Laurence, W. F. and Bierregaard Jr. R. O. (ed.). Tropical Forest Remnants - Ecology, Management and Conservation of Fragmented Communities. University of Chicago Press, Chicago. pp. 222–240.

MacArthur, R. H. & Wilson, E. O. (1963) An equilibrium theory of insular zoogeography. Evolution, 373–387.

MacArthur, R.H. & Wilson, E.O. (1967) The theory of island biogeography. Press Princeton, USA.

Malcolm, J. R. (1997). Biomass and diversity of small mammals in Amazonian forest fragments. In: Laurence, W. F. and Bierregaard Jr. R. O. (ed.). Tropical Forest Remnants - Ecology, Management and Conservation of Fragmented Communities. University of Chicago Press, Chicago. pp. 207–221.

Meiri, S. (2024). SquamBase—A database of squamate (Reptilia: Squamata) traits. Global Ecology and Biogeography.

Meyer, C. F. & Kalko, E. K. (2008). Assemblage level responses of phyllostomid bats to tropical forest fragmentation: land bridge islands as a model system. Journal of Biogeography, 35(9): 1711–1726.

Miranda, R. B., Brandão, R. A., O’Connell, K., Colli, G. R., Tonini, J. F., & Pyron, R. A. (2023). Multilocus environmental adaptation and population structure in the Cerrado gecko *Gymnodactylus amarali* (Sauria, Phyllodactylidae) from Serra da Mesa Hydroelectric Plant, Central Brazil. Frontiers in Ecology and Evolution, 11: 980777.

Oksanen, J. (2011) Vegan: community ecology package. R package ver. 2.0–2. https://CRAN.R-project.org/package=vegan.

Palmeirim, A. F., Vieira, M. V. & Peres, C. A. (2017). Non-random lizard extinctions in land-bridge Amazonian forest islands after 28 years of isolation. Biological Conservation, 214: 55–65.

Palmeirim, A. F., Benchimol, M., Vieira, M. V. & Peres, C. A. (2018). Small mammal responses to Amazonian forest islands are modulated by their forest dependence. Oecologia, 187(1): 191–204.

Palmeirim, A. F., Santos-Filho, M. & Peres, C. A. (2020). Marked decline in forest-dependent small mammals following habitat loss and fragmentation in an Amazonian deforestation frontier. PloS one, 15(3): e0230209.

Palmeirim, A. F., Farneda, F. Z., Vieira, M. V. & Peres, C. A. (2021). Forest area predicts all dimensions of small mammal and lizard diversity in Amazonian insular forest fragments. Landscape Ecology, 1-18.

Pavan, D. (2001). Considerações ecológicas sobre a fauna de sapos e lagartos de uma área do Cerrado brasileiro sob o impacto do enchimento do reservatório de Serra da Mesa. Unpublished Ms. Thesis. Universidade de São Paulo. 159 p.

Pigot, A. L., & Etienne, R. S. (2015). A new dynamic null model for phylogenetic community structure. Ecology Letters, 18(2): 153–163.

Pinheiro, J. C., & Bates, D. M. (2000). Linear mixed-effects models: basic concepts and examples. Mixed-effects models in S and S-Plus, 3–56.

Redford, K. H., & Peters, G. (1986). Notes on the biology and song of the red-legged seriema (*Cariama cristata*). Journal of Field Ornithology, 261–269.

Ricklefs, R. E.; Cochram, D. & Pianka, E. R. (1981). A morphological analysis of the structure of communities of lizards in desert habitat. Ecology, 62:1474–1483.

Sarre, S.; Smith, G. T. & Meyers, J. A. (1995) Persistence of two species of gecko (Oedura reticulata and Gehyra variegata) in remnant habitat. Biological Conservation 71:25–33.

Sarre. S. (1998) Demographics and population persistence of Gehyra variegata (Gekkonidae) following habitat fragmentation. Journal of Herpetology, 32:153–162.

Smith, G. T.; Arnold, G. W.; Sarre, S.; Abensperg-Traun, M. & Steven, D. E. (1996) The effect of habitat fragmentation and livestock grazing on animal communities in remnants of gimlet *Eucalyptus salubris* woodland in the Western Australian wheatbelt. II. Lizards. Journal of Applied Ecology, 33: 1302–1310.

Simberloff, D. S. (1974). Equilibrium theory of island biogeography and ecology. Annual Review of Ecology and Systematics, 5(1): 161–182.

Somers, K. M. (1986) Multivariate allometry and removal of size with principal components analysis. Systematic Zoology 35:359–368.

Souza, E., Lima-Santos, J., Entiauspe-Neto, O. M., dos Santos, M. M., de Moura, P. R., & Hingst-Zaher, E. (2022). Ophiophagy in Brazilian birds: a contribution from a collaborative platform of citizen science. Ornithology Research, 30(1): 15–24.

Tabachnick, B. G. & Fidell, L. S. (2001) Using Multivariate Statistics. Allyn & Bacon Inc. Boston. 966 pp.

Terborgh, J. W. & Winter, B. (1980). Some causes of extinction. In: Soulé, M. E. and Wilcox, B. A. (eds). Conservation Biology: An Evolutionary-Ecological Perspective. Sinauer Associates, Sunderland, Mass. pp. 119–133.

Terborgh, J. W., Lopez, L., Tello, J., Yu, D. & Bruni, A, R. (1997). Transitory states in relaxing ecosystems of land bridge islands. In: Laurence, W. F. and Bierregaard Jr. R. O. (ed.). Tropical Forest Remnants - Ecology, Management and Conservation of Fragmented Communities. University of Chicago Press, Chicago. pp. 256–273.

Terborgh, J., Lopez, L., Nuñez, P., Rao, M., Shahabuddin, G., Orihuela, G.,… & Balbas, L. (2001). Ecological meltdown in predator-free forest fragments. Science, 294(5548): 1923–1926.

Terborgh, J., Feeley, K., Silman, M., Nuñez, P. & Balukjian, B. (2006). Vegetation dynamics of predator free land bridge islands. Journal of Ecology, 94(2): 253–263.

Therneau T.M. (2022). coxme: Mixed Effects Cox Models. R package version 2.2-18.1, https://CRAN.R-project.org/package=coxme

Therneau T (2024). A Package for Survival Analysis in R. R package version 3.7-0, https://CRAN.R-project.org/package=survival.

Vitt, L. J. (1991). An introduction to the ecology of Cerrado lizards. Journal of Herpetology, 79–90.

Vitt, L. J., Shepard, D. B., Caldwell, J. P., Vieira, G. H. C., França, F. G. R. & Colli, G. R. (2007). Living with your food: geckos in termitaria of Cantão. Journal of Zoology, 272(3): 321–328.

Wang, Y., Zhang, J., Feeley, K. J., Jiang, P. & Ding, P. (2009). Life history traits associated with fragmentation vulnerability of lizards in the Thousand Island Lake, China. Animal Conservation, 12(4): 329–337.

Whittaker, R.H. (1972). Evolution and measurement of species diversity. Taxon, 21: 213–251.

Wilcox, B. A. (1978) Supersaturated island faunas: a species-age relationship for lizards on post-Pleistocene land-bridge islands. Science, 199(4332): 996–998.

Yahner, R. H. (1991). Dynamics of a small mammal community in a fragmented forest. American Midland Naturalist, 127:381–391.

Yu, M., Hu, G., Feeley, K. J., Wu, J. & Ding, P. (2012). Richness and composition of plants and birds on land bridge islands: Effects of island attributes and differential responses of species groups. Journal of Biogeography, 39(6): 1124–1133.

