## Supplemental Table for "How do body size and habitat fragmentation influence extinction in lizards? A long-term case study on artificial islands in the Brazilian Cerrado"

Supplementary material. Description of data sources.

| Species | Family | Mean SVL | Mean mass |
|---|---|---|---|
| **Anolis brasiliensis** | Anolidae | fieldwork | fieldwork |
| **Anolis meridionalis** | Anolidae | fieldwork | fieldwork |
| **Cercosaura ocellata** | Gymnophtalmidae | fieldwork | Meiri 2024 |
| **Colobosaura modesta** | Gymnophtalmidae | fieldwork | Meiri 2024 |
| **Micrablepharus maximiliani** | Gymnophtalmidae | fieldwork | fieldwork |
| **Iguana iguana** | Iguanidae | Meiri 2024 | Meiri 2024 |
| **Gymnodactylus amarali** | Phyllodactylidae | fieldwork | fieldwork |
| **Polychrus acutirostris** | Polychrotidae | fieldwork | fieldwork |
| **Copeoglossum nigropunctatum** | Scincidae | fieldwork | fieldwork |
| **Notomabuya freanata** | Scincidae | fieldwork | fieldwork |
| **Coleodactylus brachystoma** | Sphaerodactylidae | fieldwork | fieldwork |
| **Ameiva ameiva** | Teiidae | Person. Commun. | Meiri 2024 |
| **Ameivula ocellifera** | Teiidae | fieldwork | fieldwork |
| **Salvator merianae** | Teiidae | Meiri 2024 | Meiri 2024 |
| **Tupinambis quadrilineatus** | Teiidae | Meiri 2024 | Meiri 2024 |
| **Tropidurus montanus** | Tropiduridae | fieldwork | fieldwork |
| **Tropidurus oreadicus** | Tropiduridae | fieldwork | fieldwork |
